# A role for pH dynamics regulating transcription factor DNA binding selectivity

**DOI:** 10.1101/2024.05.21.595212

**Authors:** Kyle P. Kisor, Diego Garrido Ruiz, Matthew P. Jacobson, Diane L. Barber

## Abstract

Intracellular pH (pHi) dynamics regulates diverse cell processes such as proliferation, dysplasia, and differentiation, often mediated by the protonation state of a functionally critical histidine residue in endogenous pH sensing proteins. How pHi dynamics can directly regulate gene expression and whether transcription factors can function as pH sensors has received limited attention. We tested the prediction that transcription factors with a histidine in their DNA binding domain (DBD) that forms hydrogen bonds with nucleotides can have pH-regulated activity, which is relevant to more than 85 transcription factors in distinct families, including FOX, KLF, SOX and MITF/Myc. Focusing on FOX family transcription factors, we used unbiased SELEX-seq to identify pH-dependent DNA binding motif preferences, then confirm pH-regulated binding affinities for FOXC2, FOXM1, and FOXN1 to a canonical FkhP DNA motif that are 2.5 to 7.5 greater at pH 7.0 compared with pH 7.5. For FOXC2, we also find greater activity for an FkhP motif at lower pHi in cells and that pH-regulated binding and activity are dependent on a conserved histidine (His122) in the DBD. RNA-seq with FOXC2 also reveals pH-dependent differences in enriched promoter motifs. Our findings identify pH-regulated transcription factor-DNA binding selectivity with relevance to how pHi dynamics can regulate gene expression for myriad cell behaviours.

## Introduction

Intracellular pH (pHi) dynamics is increasingly recognized as a critical regulatory signal for many cell behaviours, including cell proliferation, migration, and differentiation. Although changes in pHi were previously viewed as a mechanism to maintain homeostasis, it is now known that changes in pHi occur and are necessary for normal cell processes such as cell cycle progression for proliferation ^1–3^, actin filament and cell-substrate adhesion remodeling for directed cell migration ^4–9^, and stem cell differentiation and lineage specification ^10–12^. Moreover, it is now established that pHi dynamics is dysregulated in human diseases, including cancer ^13–15^, diabetes ^16,17^, and neurodegeneration ^18,19^. The molecular mechanisms by which pHi dynamics regulates normal and pathological cell behaviours is often mediated by protein electrostatics and the protonation state of key residues in endogenous pH sensitive proteins termed “pH sensors” ^20,21^.

Of all amino acids that undergo protonation state changes in solution, histidine side chains exhibit the closest pKa (6.5) to physiological pH. However, the pKa of histidine, aspartic acid, glutamic acid, and buried lysine can be upshifted or downshifted depending on the protein electrostatics environment to enable titration within the cellular pH range of 7.0-7.8 ^6,22,23^.

Changes in the protonation state of key amino acids in response to small changes in pH (0.2-0.3) can significantly alter protein structure and function, including activity ^7,24–26^ and ligand binding ^5,6,27–29^ as well as stability ^30–32^ and aggregation ^33,34^. While many cell processes regulated by pHi dynamics include changes in gene expression ^3,12,35^, and although nuclear and cytosolic pH are similar ^36^, how pHi can regulate transcription factor target gene selectivity remains understudied and unclear.

Our current understanding of how transcription factors achieve DNA-binding specificity includes distinct regulatory mechanisms, such as co-factor association, post-translational modifications such as phosphorylation, and cell specific expression ^37–39^ as well as DNA architecture ^40–43^. Here, we tested the prediction that DNA binding specificity of transcription factors with a histidine in the DNA binding domain (DBD) that interacts directly with nucleotides might be regulated by intracellular pHi dynamics. Significantly, this prediction could be relevant to more than 85 transcription factors in distinct families, including FOX, KLF, SOX and MITF/Myc, which in available structures contain a histidine in the DBD that directly forms a hydrogen bond with DNA nucleotides.

We confirmed our prediction by showing pH-regulated DNA binding of three FOX family transcription factors, FOXC2, FOXM1, and FOXN1. For FOXC2 we find pHi regulated activity in cells and show the critical importance of a conserved histidine for pH-sensitive DNA binding and gene expression. Additionally, we used unbiased approaches of SELEX-seq and RNA-seq to identify different DNA motif binding profiles at lower pH (7.0-7.4) compared with higher pH (7.7-7.8). Our findings fill gaps in our current understanding not only for how pHi dynamics can tune the selectivity for target genes and regulate gene expression, but also for how transcription factors with highly similar DNA-binding domains can regulate diverse genes and functions reiteratively for developmental programs that include changes in pHi ^11,44^.

## Materials and Methods

### Amino acid sequence and structural alignment

For sequence alignment of FOX, KLF, SOX, and MITF/MYC/MAX family proteins, FASTA sequences were downloaded from UniProt and uploaded to Jalview software. For each family, sequences were aligned using ClustalO default settings. Amino acids were arbitrarily colored with red used to highlight the conserved DNA-binding histidine in each family. For structural alignment, available FOX crystal structures in complex with DNA were downloaded from the Protein Data Bank (PDB), which included FOXA2 (5X07), FOXC2 (6AKO), FOXG1 (7CBY), FOXH1 (7YZ7), FOXK2 (2C6Y), FOXL2 (7VOU), FOXM1 (3G73), FOXN1 (6EL8), FOXN3 (6NCE), FOXO1a (3CO6), FOXO3 (2UZK), FOXO4 (3L2C), FOXP2 (2AS5), and FOXP3 (7TDW). Structures were aligned using FOXC2 (green) as a reference for other FOX structures shown in complex with DNA (grey) by using PyMOL software.

### Cloning, expression, and purification

The GST-fusion protein plasmid pGEX-6P-2 was digested with BamHI and EcoRI enzymes and the DBD of FOXC2 (amino acids 72-172), FOXM1 (amino acids 222-360), and FOXN1 (amino acids 270-366) were PCR amplified with BamHI and EcoRI cloning site overhangs. FOX templates were obtained from Addgene ^45^ and FOX-DBD DNA sequences were ligated and cloned into the pGEX-6P2 using Gibson Assembly Master Mix (NEB: E2611L). Point mutants were generated with the QuikChange Lightning site-directed mutagenesis kit (Agilent: 210515) according to the manufacturer protocol. Each construct was transformed into and expressed in BL21-DE3 *E. coli* competent cells using heat shock (Thermo EC0114). For expression, cells were grown in 1L of Luria broth with ampicillin (100 µg/mL; 37°C with shaking at 225 rpm) until cells reached log-phase growth at OD_600_ = ∼0.6. Expression was induced with 1 mM isopropyl-β-D-1-thiogalatapyranoside for 6 hours at 37°C with shaking at 225 rpm. Cells were pelleted (7000g; 15 min at 4°C) and either frozen at -80°C or used directly for protein purification.

Bacterial cell pellets were resuspended in 50 ml of lysis buffer (50 mM Tris-HCl pH 8, 1 mM DTT, 5% glycerol, protease inhibitor cocktail [Roche 1183615300]). Cells in pellets were lysed by sonication on ice with 10 sec pulses of maximum setting followed by 1 min cooling period, repeated 12 times. The supernatant was clarified by centrifugation (12,000g; 30 min at 4°C) and mixed 1:1 with wash buffer (50 mM Tris, 150 mM NaCl, pH 8.0) from a GST purification kit (Pierce: 16105). 12.5 ml of lysate was loaded on pre-equilibrated 3 ml glutathione agarose spin columns, incubated end over end for 30 min at 4°C and repeated until all lysate was used. The flowthrough was collected by centrifugation (700g; 2 min at 4°C) and columns were washed with 6 ml of wash buffer three times. GST-FOX DBDs were eluted with 3 ml of wash buffer containing 10 mM reduced glutathione three times. Each fraction was collected, separated on a 10% SDS-PAGE gel, and Coomasie stained to determine molecular size and purity. Eluate fractions were pooled, divided in half, concentrated, and exchanged in two separate anisotropy buffers (20 mM Hepes, 140 mM KCl, 0.05 mM TCEP-HCl, pH 7 or 7.5) using Amicon Ultra-15 Filters with a 10 kDa molecular weight cutoff (MilliporeSigma: UFC901008). The protein concentration was determined by using a NanoDrop spectrophotometer (Thermo: ND-1000), and samples were aliquoted, flash frozen in liquid nitrogen, and stored at -80°C.

### SELEX-seq for identifying pH-dependent binding sequences

For SELEX-seq, a previously reported protocol ^46^ was used with modifications. We designed a library with a 16 base pair randomized region flanked by PCR sites GTTCAGAGTTCTACAGTCCGACGATCTGGNNNNNNNNNNNNNNNNTCGTATGCCGTCTTC TGCTTG. For each round of selective enrichment, final concentrations of 0.25 µM DNA library with 2.5 µM GST-FOXC2 were incubated for 30 min at RT in binding buffer (20 mM Hepes, 140 mM KCl, 0.05 mM TCEP-HCl) at either pH 7 or 7.8. Next, 30 µl of 50% pre-equilibrated glutathione Sepharose beads (Cytiva: 17075601) were added and samples were incubated end over end for 30 min at RT after which DNA-FOXC2-bead complexes were pelleted by centrifugation at 5000 rpm for 4 min. DNA-FOXC2-bead complexes were washed twice with 300 µl binding buffer and resuspended in 100 µl binding buffer prior to heat dissociation for 5 min at 95°C. Eluate DNA was clarified by centrifugation (10,000 rpm for 5 min) and PCR amplified with forward (SELEX_F_) and reverse (SELEX_R_) primers complementary to nucleotides flanking the randomized region (Table 1). PCR products were cleaned using the MinElute PCR purification kit (Qiagen 28004) and the selection process was repeated four times with PCR products from prior rounds used as the starting library. 5’ and 3’ adapter overhangs were added by PCR (Table 2) to the initial starting library (R_0_) with all four rounds of selection (R_1-4_) used to prepare for addition of barcoded indexes. Indexes were added by PCR (Table 2) using the Nextera IDT UD Set D (Illumina 20027213) for multiplex sequencing. Sequencing was performed at the University of California, San Francisco Genomics CoLab using a Miniseq high output 150 cycles kit for paired end reads (Illumina FC-420-1002). Samples were demultiplexed and analysis performed using the University of California, Davis Bioinformatics Core according to the protocol outlined in ^46^ using their published ‘R’ “SELEX” package which is publicly available at: (bioconductor.org/packages/release/bioc/html/SELEX.html).

**Table 1.**
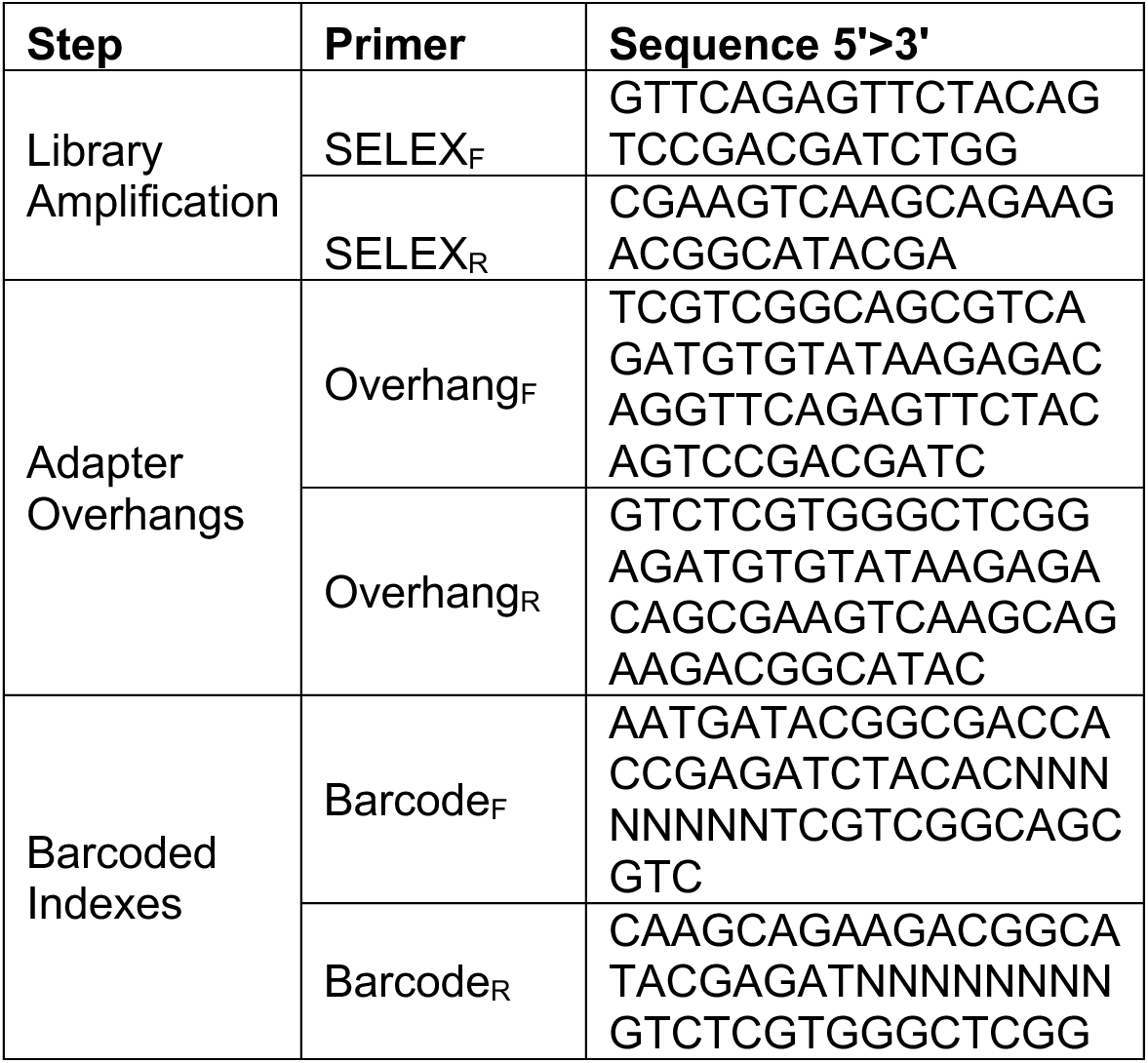
Primers used in SELEX-seq library.

**Table 2.** Indexes from Nextera IDT UD Set D used for each sample.

| Sample | Index ID | Index 5' | Index 3' |
| --- | --- | --- | --- |
| R<sub>0</sub> Initial Library | UDP0337 | TGTAAGGTGG | AAGGCCTTGG |
| R<sub>1</sub> FOXC2 pH 7.0 | UDP0338 | CAACTGCAAC | TGAACGCAAC |
| R<sub>1</sub> FOXC2 pH 7.8 | UDP0339 | ACATGAGTGA | CCGCTTAGCT |
| R<sub>2</sub> FOXC2 pH 7.0 | UDP0340 | GCAACCAGTC | CACCGAGGAA |
| R<sub>2</sub> FOXC2 pH 7.8 | UDP0341 | GAGCGACGAT | CGTATAATCA |
| R<sub>3</sub> FOXC2 pH 7.0 | UDP0342 | CGAACGCACC | ATGACAGAAC |
| R<sub>3</sub> FOXC2 pH 7.8 | UDP0343 | TCTTACGCCG | ATTCATTGCA |
| R<sub>4</sub> FOXC2 pH 7.0 | UDP0344 | AGCTGATGTC | TCATGTCCTG |
| R<sub>4</sub> FOXC2 pH 7.8 | UDP0304 | CCGCTCCGTT | TACGGCGAAG |

### Constant pH molecular dynamics

Constant pH molecular dynamics (CpHMD) simulations were performed to estimate protonation state distributions for defined titratable histidine residues in FOXC2 and FOXM1 in thermodynamic equilibrium, both as protein-DNA complexes and as solvated monomers ^47^. The pKas of histidine residues were estimated to inform how DNA-protein interaction is modulated as a function of changes in pH. All CpHMD simulations were run for 100 ns at pH 7.0, with protonation state change attempts every 100 fs.

### GST-FOX DBD anisotropy

GST-FOX DBD protein aliquots were thawed at room temperature and diluted to a final volume of 120 µl in anisotropy buffer pH 7 and 7.5 to the highest concentration of each protein (FOXC2- 50 µM, FOXM1- 21 µM, FOXN1- 50 µM, FOXC2-H122K- 9 µM, FOXC2-H122N- 97 µM, FOXM1-H287K- 21 µM) needed to reach binding saturation, which was empirically determined. Protein was serially diluted ten times to a volume of 60 µL and 10 µl of 6’FAM labeled duplex FkhP (ccATAAACAac) (IDT) was added to each protein dilution and one blank for a final concentration of 7.5 nM DNA and 70 µl reaction volume. PCR strip tubes were capped and incubated at RT in the dark for 30 min. Following incubation, 20 µl of each dilution and blank was loaded in triplicate to a 384-well black plate (Greiner: 784076) using a multichannel pipette. Fluorescence anisotropy measurements were made using a SpectraMax M5 plate reader (Molecular Devices). Sigmoidal curve fits were generated using GraphPad Prism and association constants determined using Mathematica software to solve for K_D_ in equation *A_obs_* = *A*_0_ + (Δ*A* ∗ *T*)/(*K_D_* + *T*) where A_obs_ is observed anisotropy, A_0_ is anisotropy of initial unbound probe, ΔA is difference in anisotropy between unbound and fully bound populations, and T is concentration of titrant protein. K_A_ was determined as (1/K_D_). Data are expressed as averages of at least three independent measurements from two to three separate protein preparations and error bars are ± s.e.m. Binding affinities of FOX wild type (WT) proteins were analysed by two-tailed unpaired Student’s t-test and with a significance level of p<0.05. Comparison of WT FOXC2 and FOXM1 to His mutants was analysed by Tukey-Kramer HSD with a significance level of p<0.05.

### GST-FOX DBD pH titration

Anisotropy buffer was prepared in increments of 0.2 pH units from pH 6.8-8.2 and 7.5 nM final concentration of labeled FkhP oligo was added to each. Either 2.3 µM final concentration GST- FOXC2 or buffer alone was added to a final reaction volume of 70 µl. Reactions were incubated and measured as described above. Data are represented as average anisotropy values of [(FOXC2+DNA) – (DNA alone)] set relative to maximal binding at pH 6.8 ± s.e.m.

### Cell Culture and luciferase assay

For determinations with EIPA to lower pHi, MDA-MB-436 cells obtained from ATCC and commercially authenticated (IDEXX BioAnalytics) were maintained in atmospheric conditions at 37°C in Leibovitz’s L-15 medium (Cytiva: SH30525.01) supplemented with insulin (10µg/ml), Penicillin/Streptomycin (100U/ml each), and 10% FBS. For determinations with NHE1-silenced conditions, MDA-MB-436 cells were maintained at 37°C and 5% CO_2_ in RPMI 1640 (Gibco:11875085) supplemented with Penicillin/Streptomycin (100U/mL each), and 10% FBS. For all luciferase assays, 3.5 x 10^5^ MDA-MB-436 cells were plated in 6-well plates, grown overnight to 80% confluency and then transfected with a total of 1 µg DNA using lipofectamine 3000 (Invitrogen L3000001) according to manufacture protocol. Cells received either 500 ng 6x-FkhP (6x-DBE) obtained from Addgene ^45^ or 500 ng 6x-FkhP with 500 ng pCS2 Flag-FOXC2 WT from Addgene ^45^ or mutant FOXC2 generated as described above. Each transfection also included control pRL-TK renillla plasmid obtained from the L. Selleri Lab (University of California San Francisco) at a ratio of 1:10 of reporter plasmid (50 ng). At 8 hours after transfection, cells were washed once with PBS and growth medium added in the absence or presence of 10 µM EIPA. Cells were then maintained for an additional 40 h and collected for Dual-Luciferase assays (Promega: E2920). In brief, cells were washed with PBS and lysed in 500 µl of Dual-Glo luciferase buffer with shaking on a nutator for 10 min at 4°C. Lysates were collected in microfuge tubes and clarified by centrifugation for 5 min at 13,000 rpm at RT. From supernatants, 100µl was loaded in quadruplicate in separate wells in a 96-well opaque white plate (Costar: 3917). Luciferase signal was read on a SpectraMax M5 plate reader and Dual-Glo Stop & Glo Buffer was then added to quench the luciferase signal and activate renilla for 10 min. The renilla signal was read and the Luciferase/Renilla ratio was normalized to control with 6x- FkhP + WT FOXC2. Data were analysed by Tukey-Kramer HSD with a significance level of p<0.05.

### pHi determinations

For imaging cytosolic and nuclear pH, MDA-MB-436 cells plated on 35 mm glass bottom MatTek dishes (MatTek Corporation P35G-1.5-10-C) for 48 h were washed 2X and incubated for 15 min in pHi buffer (110 mM NaCl, 5 mM KCl, 10 mM glucose, 25 mM NaHCO_3_, 1 mM KPO_4_, 1 mM MgSO_4_, 2 mM CaCl_2_, pH 7.4) containing 10 mM of the dual emission pH-sensitive dye 5-(and-6)-carboxylic acid, acetoxymethyl ester (SNARF) as previously described ^48^. For measurements with EIPA, a final concentration of 10 mM was included in cell medium 24 h before measurements and in all wash and dye-loading buffers. After dye loading and washing 2X in pHi buffer, ratiometric determinations at Ex 490 were made at pH-sensitive and - insensitive emissions of 580 nm and 640 nm, respectively. Dye ratios with imaging and as described below for cell populations were calibrated to pHi values by incubating cells at the end of each determination sequentially for 5 min each with a Na^+^-free, K^+^ buffer containing the ionophore nigericin at pH 7.5 and then at pH 6.6 to equilibrate intracellular and extracellular pH, as previously described ^48,49^ SNARF images were obtained using a Plan Apo 40 0.95 NA objective on an inverted Nikon spinning disc microscope system (Nikon Eclipse TE2000 Perfect Focus System; Nikon Instruments; Nikon Instruments) equipped with a CoolSnap HQ2 cooled charge-coupled camera (Photometrics) and camera-triggered electronic shutters controlled with NIS-Elements Imaging Software (Nikon).

For CRISPR/Cas9 knock out of NHE1 (NHE1 KO) in MDA-MB-436 cell, we used a protocol we recently described with guide RNA targeting the first exon of the NHE1 gene SLC9A1, including fluorescence activated sorting to select for cells transiently expressing Cas9- GFP ^11^. For measuring pHi in cell populations, parental MDA-MB-436 cells in the absence or presence of EIPA and NHE1 KO MDA-MB-436 cells plated in 24-well dishes for 48 h were washed 2X and incubated for 15 min in pHi buffer containing 1 mM of the dual excitation pH-sensitive dye 2’,7’-Bis-(2-Carboxyethyl)-5-(and-6)-carboxyfluorescein, acetoxymethyl ester (BCECF) as previously described ^49^. After dye loading and washing 2X in pHi buffer, ratiometric determinations were made at Em 535 nm and pH-sensitive and -insensitive excitations of 490 nm and 440 nm, respectively, using a SpectraMax M5 plate reader^47,48^. To confirm loss of NHE1 activity with EIPA or with NHE1 KO, in a nominally HCO_3_-free Hepes buffer (145 mM NaCl, 5 mM KCl, 10 mM glucose, 25 mM Hepes, 1 mM MgSO4, 1 mM KPO4, 2 mM CaCl2, pH 7.4) the rate of H+ efflux was determined by measuring the time-dependent pHi recovery from an NH_4_Cl-induced acid load as previously described ^49^.

### Library preparation, RNA-seq, and analysis

MDA-MB-436 cells were maintained as described for luciferase assays and plated at 3.5 x 10^5^ control or NHE1-KO cells per well in 6 WP and grown overnight to 80% confluency prior to transfection. Cells were either untransfected or transfected with 1 µg of FOXC2-WT, -H122K, or -H122N with medium changed 8 h after transfection. Cells were incubated for 48 h post transfection and RNA was extracted with the RNeasy Mini kit (Qiagen, 74104) according to manufacturer instructions with sample concentrations assessed via NanoDrop. Samples were frozen at -80°C and sent to Novogene Co. Ltd (USA) for library RNA sample quality assessment, library construction, RNA-sequencing on an illumina NovaSeq 6000, read mapping, and statistical analysis of differentially expressed genes. Data are from three paired biological replicates of each condition. Differentially expressed genes in each condition were analysed using InteractiVenn web tool ^50^. Volcano plot analysis was performed with a custom Python script with a significance value cutoff of (qval < -log_10_ (0.05)). Select gene fold change between controls and NHE1-silenced cells for untransfected, FOXC2-WT, -H122K, or -H122N was generated with GraphPad Prism. Promoter elements from gene lists in the human hg19 genome at -2000 to +100 base pairs from the transcription start site were analysed using the HOMER FindMotifs script. The most common promoter elements enriched at high vs low pH were compared using the DiffLogo ^51^ R package with statistical analysis of position weight matrix difference performed with Kullback-Leibler divergence.

## Results

### Predicted pH-sensing by transcription factors with a conserved histidine in the DBD

We previously described the design principles of several pH-sensors that are regulated by titration of a histidine within the cellular pH range of 7.0 to 7.8 ^5–7,26,30,52^. In asking whether some transcription factors might function as pH sensors we searched for conserved histidine residues important for DNA binding across transcription factor families. We first performed sequence alignments of major transcription factor family members expressed in humans. We find that all FOX family members contain a histidine in a highly conserved N(S/A)IRH motif within Helix 3 of the DBD (**Fig. 1A**). In all available crystal structures of FOX transcription factors in complex with DNA, the conserved histidine aligns in the major groove of DNA and forms a hydrogen bond with nucleotides. Further, we used a structural overlay to show that the position of the side chain of conserved histidine residues is spatially conserved (**Fig. 1E**). We also find that all members of the KLF transcription factor family, all members of the SOX transcription factor family except Sry and SOX30, and all members of the MITF/Myc/Max family except AP4 contain a conserved histidine in the DBD, which in available structures in complex with DNA forms a hydrogen bond with nucleotides (**Fig. 1B-D, Supplementary Figure S1A, B,C)**. Additionally, a hydrogen bond between a histidine in the DBD and DNA nucleotides is reported for the ETS transcription factor ETV6 ^53^, the STAT transcription factor STAT6 ^54^, and ARNT, the DNA-binding subunit of the HIF1 complex ^55,56^ (**Supplementary Figure S1D-F)**. Hence, at least 85 transcription factors in diverse families contain a histidine in the DBD that in available structures forms a hydrogen bond with nucleotides.

**Figure 1.**
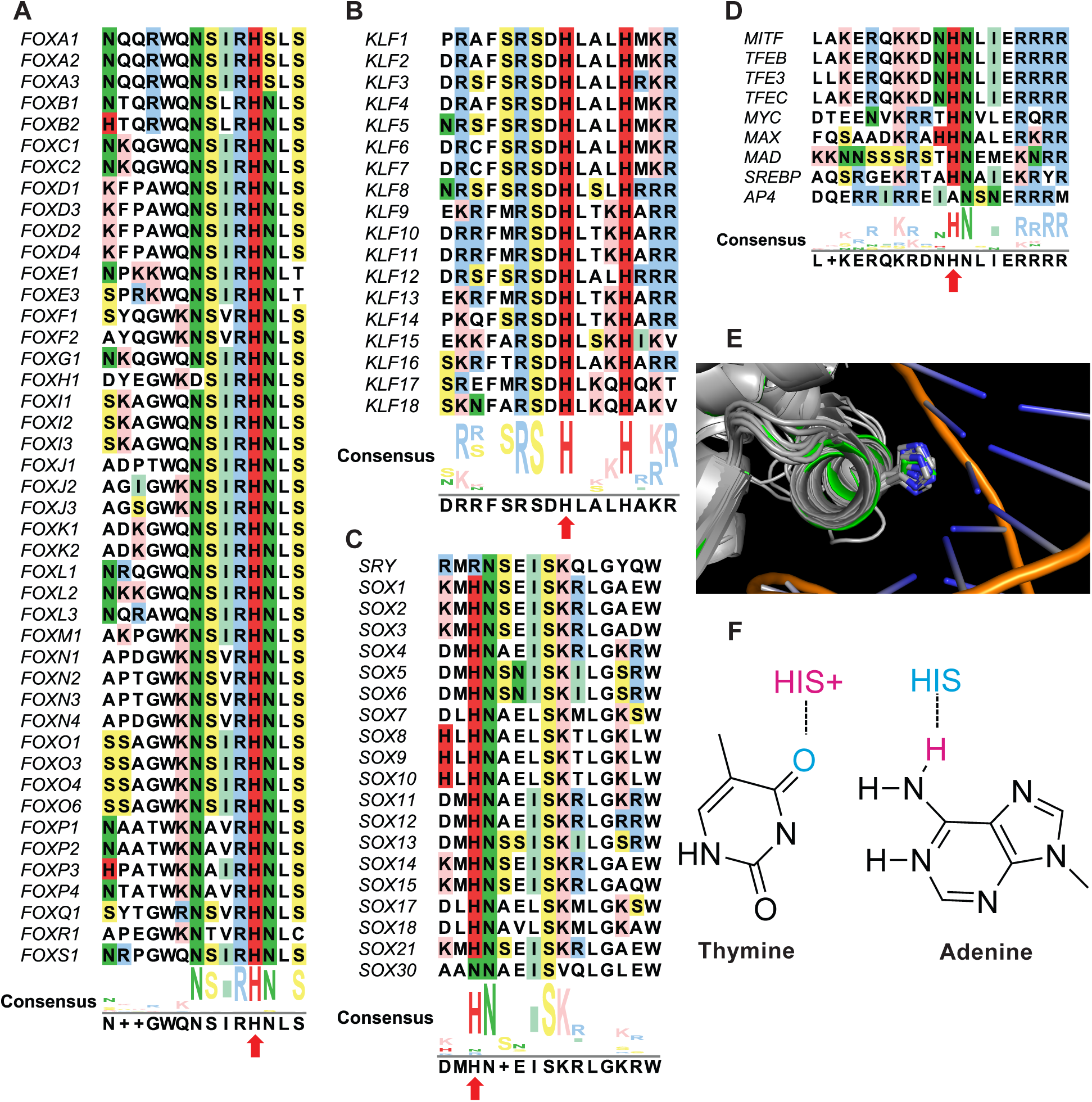
FOX family proteins with predicted pH-regulated DNA binding. A-D. Sequence alignment of Helix H3 of the DBD of FOX (A), KLF (B), SOX (C), and MITF/MYC/MAX (D) family members with conserved histidine highlighted in red and denoted by red arrows. E. Alignment of all available crystal structures of FOX proteins in complex with DNA highlighting spatial alignment the side chain of the conserved histidine. F. Proposed model for pH-regulated transcription factor-DNA binding.

Given the conservation of a histidine in the DBD of many transcription factors that forms a hydrogen bond with nucleotides, coupled with the well-established ability of histidine to titrate within the cellular pH range, we hypothesized that a pH-dependent titration of the conserved histidine might regulate DNA binding specificity through hydrogen bonding preferences with nucleotides. For example, a protonated histidine may serve as hydrogen bond donor for a hydrogen bond accepting thymine. However, when deprotonated at higher pH a deprotonated histidine may serve as a hydrogen bond acceptor for an adenine donor (**Fig. 1F**). Guanine and cytosine can be either a hydrogen bond donor or acceptor depending on the orientation of the nucleotide relative to the protein binding residue.

### SELEX-seq reveals pH-dependent binding preferences of FOXC2 *in vitro*

To test our hypothesis, we used SELEX-seq as an unbiased approach to identify pH-dependent DNA binding motifs from a randomized library (**Fig. 2A**), using a purified recombinant FOXC2 DBD expressed as a GST fusion protein (**Supplementary Figure S2**) at pH 7 and pH 7.8. We chose using FOXC2 because it is the only FOX family member with a crystal structure shown in complex with DNA that contains a single histidine (His122) in the DBD (Supplementary Figure S1). Additionally, we limited our screen to pH 7 and 7.8 as representative pH values near the physiological limits of a cell ^57,58^. After four rounds of selection at pH 7 we find that the most enriched sequence is the canonical Forkhead Primary (FkhP) motif ATAAACA (**Fig. 2B**). In contrast, at pH 7.8 the most enriched sequence is a known FOX alternative consensus motif (FHL) ^59^ GACGC and is different than at pH 7 (**Fig. 2B**). These data suggest there are pH-dependent preferences in known FOX DNA binding motifs for at least FOXC2.

**Figure 2.**
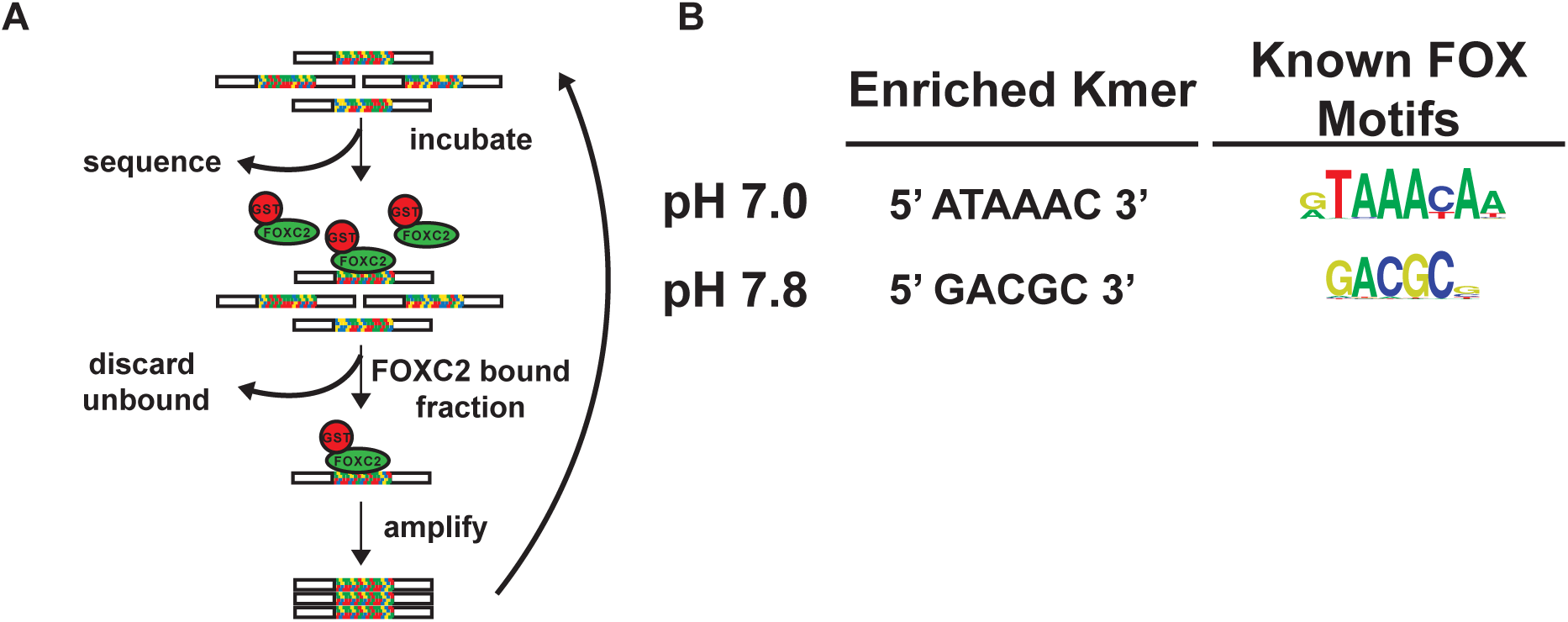
SELEX reveals pH-dependent differences in DNA binding sequences for FOXC2. A. Design of SELEX assay. B. Most enriched sequences in SELEX-seq after four rounds of enrichment, each at pH 7.0 or 7.8 compared to known FOX consensus motifs from Nakawa et al. ^59^.

### FOX family transcription factors have higher affinity for an FkhP motif at lower pH

We tested the predicted protonation state of FOXC2-His122 when bound to the SELEX identified FkhP sequence *in silico* using CpHMD. Our data suggest that when FOXC2 is bound to the FkhP sequence, His122 is predicted to be doubly protonated in 91% of the simulation. Single protonated populations at the delta or epsilon nitrogen for FOXC2 in complex with DNA are observed only in 1% and 8% of the simulation, respectively. In contrast, when FOXC2 is in the unbound state His122 is predicted to be preferentially singularly protonated at the delta or epsilon nitrogen, 81% and 14% of the simulation respectively, while only doubly protonated in 5% (**Fig. 3A**). Together, these data suggest that FOXC2-His122 is a putative pH-sensing residue predicted to bind the canonical FkhP sequence when protonated at low pH in agreement with findings from SELEX-seq.

**Figure 3.**
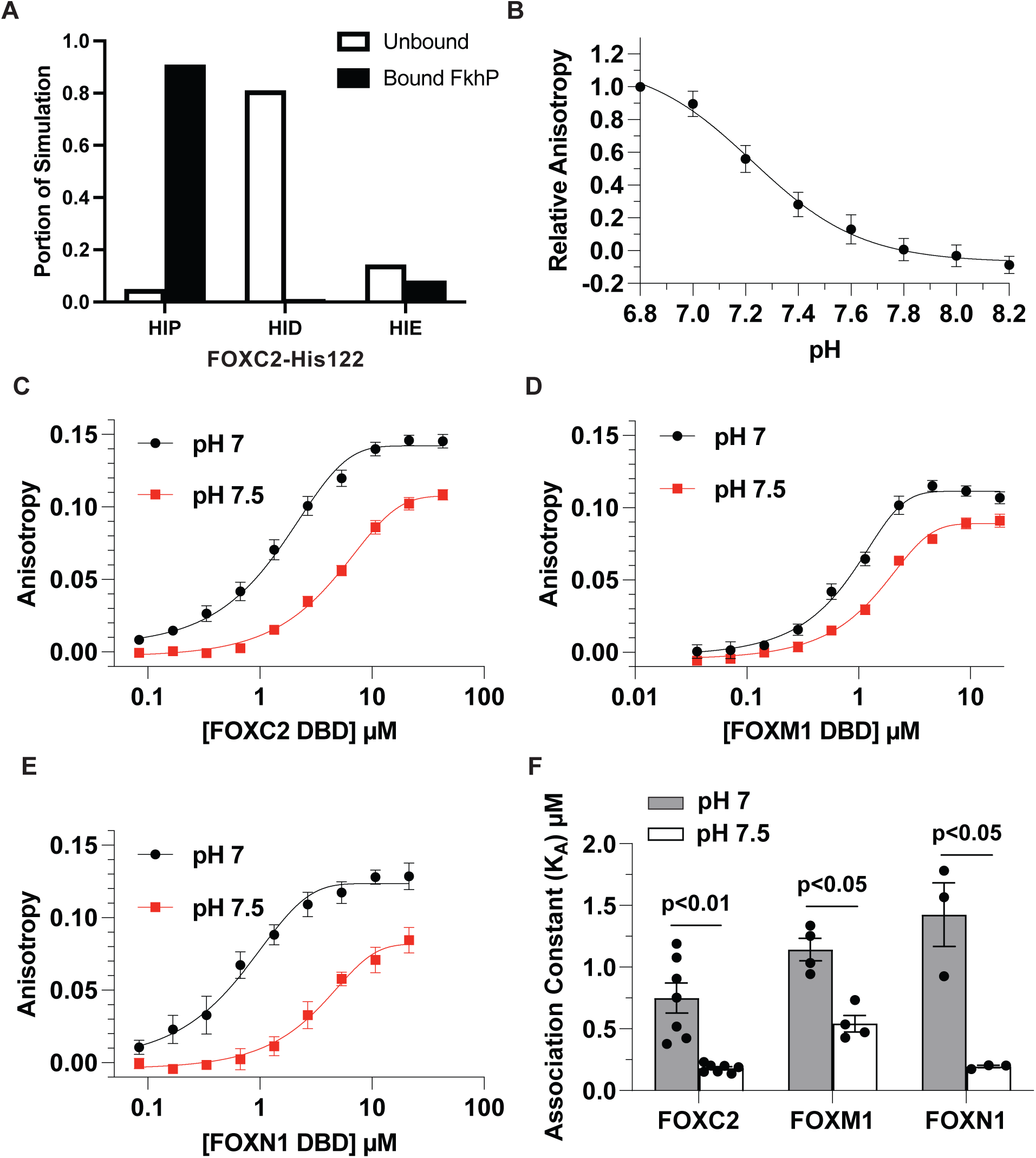
Binding of FOX family proteins to an FkhP sequence is pH-dependent with higher affinity at lower pH. A. CpHMD simulations with FOXC2-His122 double protonated (HIP), protonated at delta nitrogen (HID), or protonated at epsilon nitrogen (HIE). Distribution of protonation states in simulations of unbound FOXC2 is in white bars and bound to an FkhP in black bars for each protonation state. B. pH titration of recombinant FOXC2 binding to an FkhP sequence. C-E. Binding curves at pH 7.0 and 7.5 for FOXC2 (C), FOXM1 (D), and FOXN1 (E). F. Association constants for FOXC2, FOXM1 and FOXN1 binding to an FkhP sequence at pH 7.0 and 7.5 calculated from binding curves. Data are means ± s.e.m. of at least three separate measurements with two-three independent recombinant protein preparations. Statistical analysis by Student’s paired t-test.

We next asked whether binding of FOX family proteins to the canonical FkhP sequence has higher affinity at low pH *in vitro,* as predicted by both SELEX and CpHMD simulations. We first tested this prediction for FOXC2 by using fluorescence anisotropy with the previously described GST-FOXC2-DBD and a 5’ 6-FAM labeled FkhP sequence. A pH titration of FOXC2 at sub-saturation protein concentration of 2.3 µM reveals that the overall affinity for the FkhP sequence decreases linearly within the cellular pH range between pH 7.0 to 7.8 with no observable change in binding above pH 7.8 at this protein concentration (**Fig. 3B**). We next determined the association constant (K_A_) for FOXC2 to the FkhP sequence at pH 7 and 7.5 and find that the binding affinity of FOXC2 for the FkhP sequence is significantly greater at pH 7.0 (K_A_ of 0.75 ± 0.12 µM) compared with 7.5 (0.18 ± 0.02 µM) (**Fig. 2C, F**). We also asked whether other FOX family members have higher affinity binding to the FkhP sequence at lower pH. Using GST fusions of DBD sequences (**Supplementary Figure S2B, C)**, we find higher affinity binding for FOXM1 at pH 7.0 (K_A_ 1.1 ± 0.11 µM) compared with pH 7.5 (0.54 ± 0.08 µM) (**Fig. 3D, F**) and for FOXN1 at pH 7 (K_A_ 1.4 ± 0.31 µM) compared with pH 7.5 (0.19 ± 0.01 µM) (**Fig. 3E, F**). These findings indicate that binding of three FOX family members, FOXC2, FOXM1, and FOXN1, to the FkhP sequence is pH-dependent, with higher affinity binding at pH 7.0 compared with pH 7.5.

### His122 of FOXC2 is necessary for pH-dependent binding to the FkhP sequence

To determine the significance of the conserved histidine for pH-regulated binding to the FkhP sequence we focused on FOXC2-His122 because it is the only histidine in the FOXC2 DBD. We used site-directed mutagenesis, first testing a His122Lys substitution with the prediction that lysine with a pKa of > 10 in solution would be constitutively charged within the cellular pH range and a hydrogen bond donor analogous to a protonated histidine. We find that FOXC2-H122K has strong relative binding affinity for the FkhP sequence between pH 6.8-8 (**Fig. 4A**). Further, when we performed a FOXC2-H122K protein titration, we find that the affinity at pH 7.0 (K_A_ 0.86 ± 0.15 µM) is similar to FOXC2-WT at pH 7.0 but pH-independent with no significant difference at pH 7.5 (K_A_ 0.91 ± 0.08 µM) (**Fig. 4B, D**). We also tested a His122Asn substitution, with the prediction that an asparagine would mimic a deprotonated histidine and have lower affinity and pH-independent binding to the FkhP sequence. Moreover, a naturally occurring FOXC2-H122N mutation is reported in lung cancers ^60^. Our data confirm that the binding affinity of a FOXC2-H122N DBD is lower compared with WT at pH 7.0 and similar to WT at pH 7.5, but also pH-independent with no difference in affinity at pH 7.0 (K_A_ 0.14 ± 0.03 µM) compared with pH 7.5 (K_A_ 0.10 ± 0.01 µM) (**Fig. 4C, D**).

**Figure 4.**
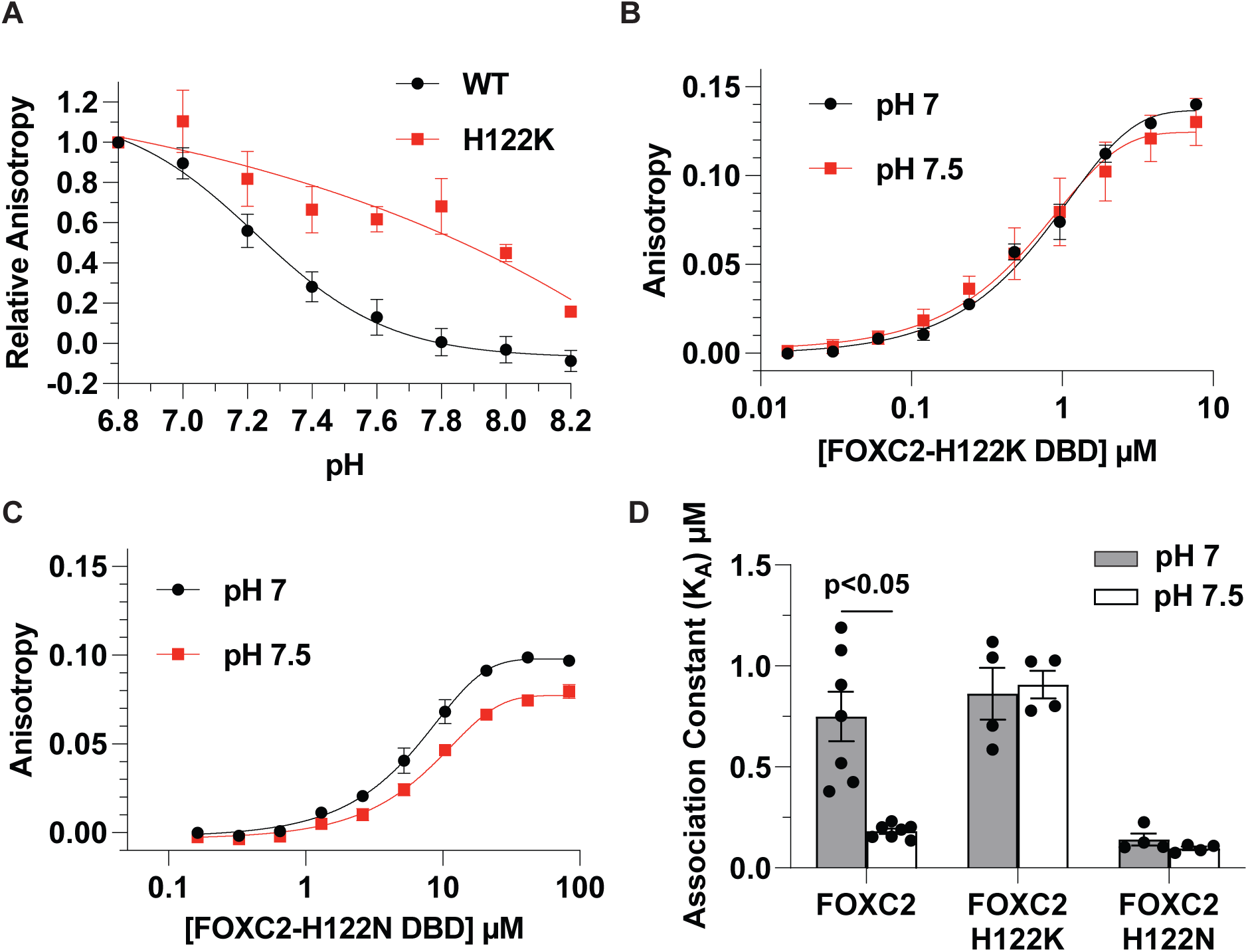
pH-dependent binding of FOXC2 to an FkhP sequence is dependent on His122. A. pH titration of FOXC2-WT and -H122K for binding an FkhP sequence. B, C. Binding curves of predicted pH-independent mutants FOXC2-H122K (B) and -H122N (C). D. Association constants for FOXC2-WT, -H122K and -H122N calculated from binding curves. Data are means ± s.e.m. of at least three separate measurements with two-three independent protein preparations and statistical analysis by Tukey-Kramer HSD test.

We also tested the importance of the conserved His287 in FOXM1 for pH-dependent DNA binding by using a His287Lys substitution. The affinities for FOXM1-H287K at pH 7 (K_A_ 4.1 ± 0.81 µM) and pH 7.5 (K_A_ 1.8 ± 0.27 µM) are both greater compared with FOXM1-WT at these pH values; however, binding is still pH-dependent (**Supplementary Figure S3A, B).** To determine why FOXM1-His287 is the not the sole determinant for pH-dependent binding, we used CpHMD simulations to sample five histidine residues in the FOXM1-DBD. Our results predict that the protonation state of His269 (blue), His275 (cyan), and His311 (green) does not affect DNA binding. In contrast, both His287 (orange) and the proximal His292 (yellow) are predicted to prefer the DNA bound state when protonated and unbound state when neutral (**Figure S3**). Together, these data indicate that His122 is necessary for pH-dependent binding of FOXC2, but in FOX family proteins with more than one histidine in the DBD, the conserved histidine may contribute to but not exclusively confer pH-dependent DNA binding.

### FOXC2 has pHi-dependent activity in cells with greater activity for an FkhP reporter at lower pHi

We next asked whether pHi regulates FOXC2 activity for an FkhP sequence in cells, using a luciferase assay with MDA-MB-436 clonal human breast cancer cells. Imaging MDA-MB-436 cells loaded with the pH-sensitive dye SNARF, we find a relatively high pHi of 7.62 ± 0.21, which is common in cancer cells ^13,14^ that is decreased to pHi 7.42 ± 0.04 in the presence of 5-(N-Ethyl-N-isopropyl)-Amiloride (EIPA) (10 µM, 24 h), a selective pharmacological inhibitor of the plasma membrane H+ extruder NHE1 (**Fig. 5A, Fig. S4A**). Imaging also revealed that pHi and nuclear pH are similar in control cells and cells treated with EIPA (**Fig. 5A**). In MDA-MB-436 cells transfected with a 6x-FkhP repeat luciferase reporter containing a minimal promoter (minP) in the absence of heterologous expressed FOXC2, there is minimal basal reporter activity with no significant difference between controls and cells treated with 10 µM of EIPA (**Fig. 5B**). In cells transfected with FOXC2-WT, however, there is a significantly greater relative reporter signal in the presence compared with the absence (Control) of EIPA (**Fig. 5B**). We also tested activity of mutant FOXC2-H122K and FOXC2-H122N in MDA-MB-436 cells with the 6x-FkhP repeat luciferase reporter. We find that both mutant proteins have measured activity, but it is pH-independent (**Fig. 5B**), consistent with their pH-independent binding affinities for the FkhP sequence determined *in vitro* by fluorescence anisotropy (**Fig. 4B-D**). Although we find pH-independent luciferase activity in cells expressing FOXC2-H122K and FOXC2-H122N, the overall activity of FOXC2-H122K was lower than WT and lower than expected based on the observed high affinity *in vitro*. This difference may be due to changes to tertiary structure of the full-length mutant protein compared with the shorter DBD used for *in vitro* binding measurements.

**Figure 5.**
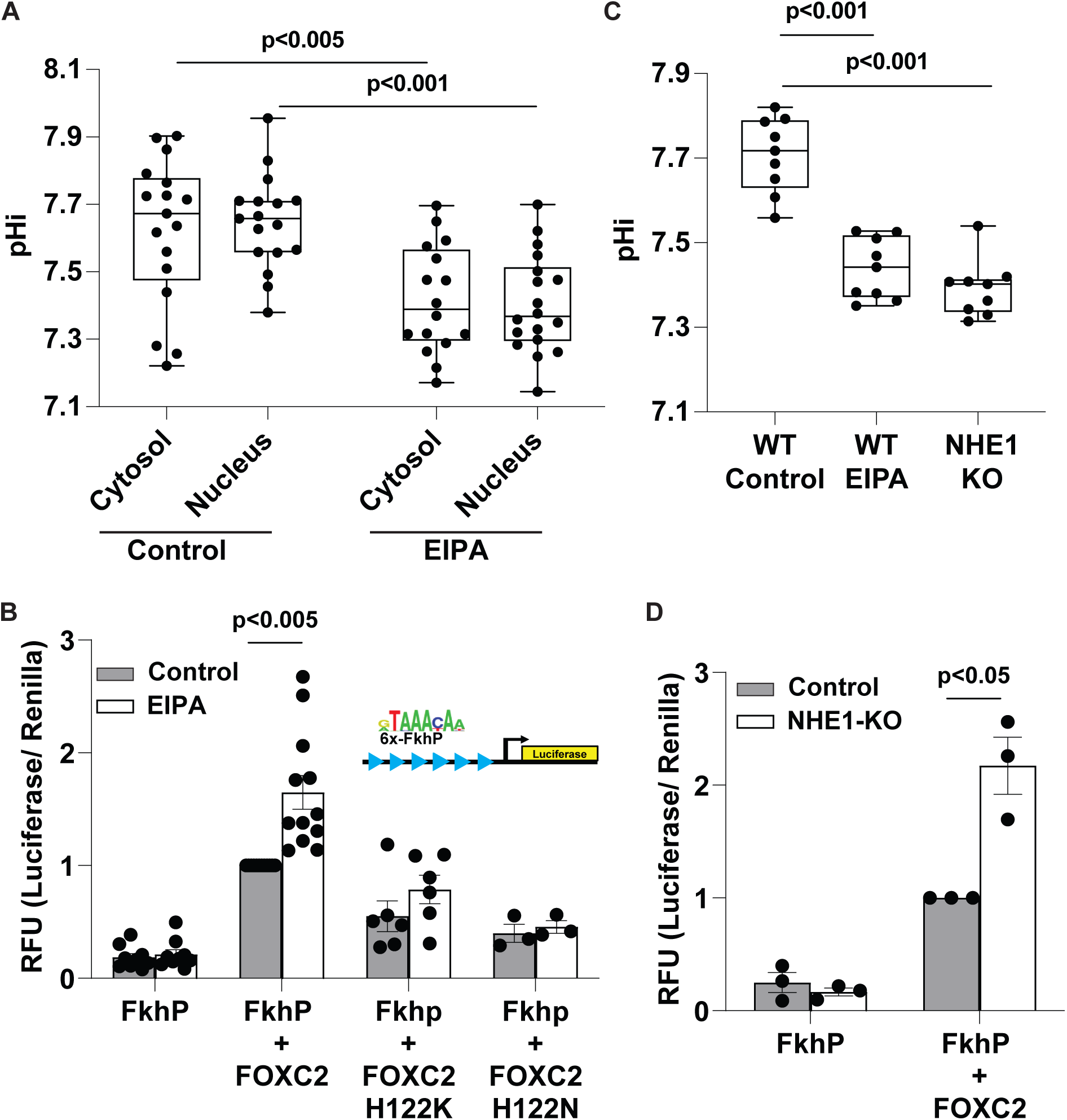
FOXC2 has pH-dependent transcriptional activity in cells determined by His122. A. The cytosolic and nuclear pH of MDA-MB-436 cells in the absence (Control) and presence of EIPA (10 mM, 24h). Box plots show median, first and third quartile, with whiskers extending to observations within 1.5 times the interquartile range, and individual data points representing an individual cell and including data from three separate cell preparations. Statistical analysis by Tukey-Kramer HSD test. B. Activity of FOXC2-WT, -H122K, and -H122N determined using a 6X-FkhP luciferase reporter (insert) in MDA-MB-436 cells in the absence (Control; pHi ∼7.65) and presence of EIPA (pHi ∼7.4) cells. Data are means ± s.e.m of three to four separate cell preparations with statistical analysis by Tukey-Kramer HSD test. C. pHi of MDA-MB-436 cell populations, including WT in the absence (Control) and presence of EIPA (10 mM, 24h) and CRISPR/Cas9-generated deletion of NHE1 (NHE1 KO). Box plots and statistical analysis as described in (A) from 3 separate cell preparations. Each data point represents the pHi value per well as a population of cells. D. Activity of wild-type FOXC2 determined using a 6X-FkhP luciferase reporter in MDA-MB-436 cells, including WT in the controls and NHE1-silenced cells. Data are means ± s.e.m of three separate cell preparations with statistical analysis by Student’s paired t-test.

To lower pHi, we added EIPA 8 h after transfecting with FOXC2 and the luciferase reporter. As an alternative approach to eliminate a time delay, we generated MDA-MB-436 cells with CRISPR/Cas9 NHE1 KO. Using cell populations loaded with the pH-sensitive dye BCECF, we see a lower pHi of 7.39 ± 0.03 with NHE1 KO compared with a pHi of 7.71 ± 0.03 in control parental cells (**Fig. 5C**). Additionally, we confirmed loss of NHE1 activity in WT MDA-MB-436 cells treated with EIPA and with NHE1 KO by showing no pHi recovery in a nominally HCO_3_-free Hepes buffer from an NH_4_Cl-induced acid load compared with pHi recovery of WT cells (Fig. S4B), which is an index of NHE1-dependent H+ extrusion. With NHE1 KO, like with EIPA-treated WT cells, co-expression of the reporter and FOXC2 results in a >2-fold increase in activity compared with activity in parental controls (**Fig. 5D**) and also in the absence of FOXC2 the reporter signal is minimal and pH-independent (**Fig. 5D**). Further, we confirmed these results are not due to differential FOXC2 expression in WT compared with NHE1 KO cells (**Fig. S4C**). Together, these data using two different approaches to lower pHi, pharmacological and genetic, show that FOXC2 activity for an FkhP reporter in cells is regulated by pHi, with greater activity at lower pHi likely mediated by His122.

### RNA-seq identifies pH-dependent differences in gene expression and suggested binding motifs

After confirming our first prediction of higher affinity binding to thymine in the canonical FkhP motif at lower pH, we were unable to observe measurable binding to confirm high pH high affinity binding of FOXC2 to several AGC-rich sequences including the SELEX identified FHL sequence (or a 2x repeat) nor other published FOXC2 binding sites in the promoters of p120 (ccAGAAAATGTATGac) and N-cadherin (ccACAAATAac) ^61,62^. Given the difficulties of validating our high pH high affinity SELEX hits *in vitro*, we asked whether we can identify high affinity motifs at higher pH by determining differences in gene expression and enriched promoter elements dependent on both pHi and FOXC2-His122. To establish genes that are both pH and FOXC2-His122 dependent we performed bulk RNA-seq in control (pH 7.71) and NHE1 KO (pH 7.42) MDA-MB-436 cells that were either untransfected, transiently expressing recombinant FOXC2-WT, or transiently expressing pH-independent mutants FOXC2-H122K or FOXC2-H122N. We find that in cells transfected with FOXC2-WT there are 2174 differentially expressed genes (DEGs) between WT and NHE1 KO cells. However, we find that only 231 of these DEGs (green circle) are both pH and FOXC2-His122 dependent due to lack of differential expression of these genes in untransfected or with pH-independent mutants (**Fig. 6A**). Further, we determined which of the pH and FOXC2-His122 dependent DEGs were enriched at a higher pH of 7.71 (controls) compared with lower pHi of 7.42 (NHE1 KO). Of the 231 DEGs, we find 113 are significantly enriched in controls at higher pHi while 118 are enriched at lower pHi in NHE1 KO cells (**Fig. 6B).** Selected genes enriched at higher pHi included UGDH, ABHD12, FGD4, ENFA2, and FGB, which promote epithelial to mesenchymal transition, ferroptosis resistance, cell migration and proliferation, and angiogenesis (**Fig. 6C**) ^63–70^. In contrast, genes enriched at lower pHi included SPEN, EP400, KMT2D, BTK, and GYPC, which promote tumor suppression, senescence, DNA-damage response, and apoptosis (**Fig. 6D**) ^71–84^. Together these results highlight there are indeed pHi- and FOXC2-His122-dependent differences in gene expression.

**Figure 6.**
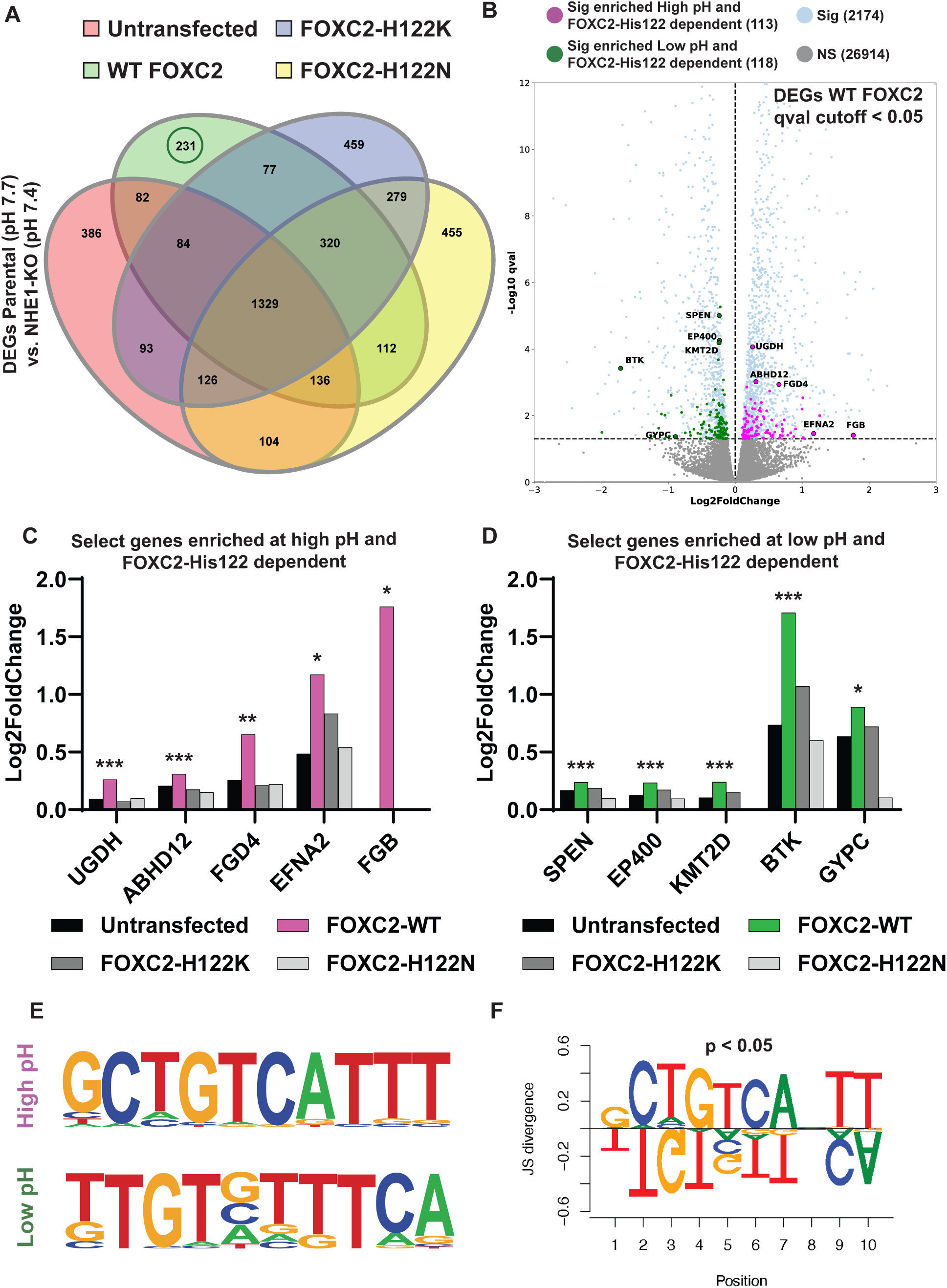
RNA-seq reveals pH-regulated and FOXC2-His122-dependent differences in gene expression and enriched motifs. A. Differentially expressed gene sets between control (pH 7.7) and NHE1 KO (pH 7.4) MDA-MB-436 cells in untransfected (red), overexpressing FOXC2-WT (green), FOXC2-H122K (blue), or FOXC2-H122N (yellow). B. Volcano plots of DEGs (qval < 0.05) in control compared with NHE1 KO MDA-MB-436 cells overexpressing FOXC2-WT. Each dot represents a single gene with pH- and FOXC2-His122 dependent genes enriched at high (magenta) vs low (green) pH and all other DEGs (blue). C, D. Expression of enriched genes at higher pH (C) or lower pH (D) with asterisks indicating qval where ***, **, and * are q < 0.001, 0.005, and 0.05, respectively. E. Common promoter elements in genes enriched at high and low pH identified by HOMER. F. Position weight matrix comparison of promoter elements.

To determine whether there are common promoter elements of the DEGs enriched at higher vs lower pHi are different, we used HOMER, a functional enrichment analysis for motif discovery, which reveals different motifs at higher compared with lower pHi (**Fig. 6E,F**). Notably, in the second nucleotide position where FOXC2-His122 is predicted to make a hydrogen bond, motifs enriched at higher pHi contain either a cytosine or adenine while motifs enriched at lower pHi contain an invariant thymine as our model predicts (**Fig. 6F**). However, using fluorescence anisotropy measurements we do not observe significant binding of FOXC2 to these HOMER identified motifs at either pH 7 or 7.5, suggesting that in cells other regulatory elements may be required for high affinity binding. Although we could not confirm with *in vitro* binding a motif with higher affinity at higher pH for FOXC2, different motif preferences at lower and higher pH suggested by SELEX-seq and RNA-seq warrants further investigation.

## Discussion

We report on an underappreciated role of pH dynamics in regulating transcription factor-DNA binding selectivity. The concept is relevant to transcription factors with a histidine in the DBD that forms hydrogen bonds directly with DNA nucleotides and could apply to at least 85 transcription factors across multiple families, as shown in Fig. 1A and Fig. S1. We do not predict that pHi dynamics functions as binary switch for DNA binding preference but rather acts as a coincidence regulator with other established mechanisms such as co-factor association, post-translational modifications like phosphorylation, and DNA accessibility through epigenetic modifications ^85^. For the latter, select pH-sensing epigenetic readers and writers were recently shown to regulate chromatin accessibility ^86,87^.

We confirm that three FOX family transcription factors, FOXC2, FOXM1, and FOXN1, bind the canonical FkhP DNA sequence (ATAAACA) with higher affinity at lower pH *in vitro* and for FOXC2 also in cells. Additionally, for FOXC2, pH-dependent DNA binding and activity are conferred by a conserved His122. These data are consistent with our prediction that a protonated histidine at lower pH acts as a hydrogen bond donor for thymine, which is generally a hydrogen bond acceptor. Two previous reports with other FOX family members also indicate pH-regulated functions. Using FOXP2, Blane and Fanucchi ^88^ show that the protonation state of the conserved histidine in the DBD regulates protein tertiary shape and DNA-binding affinity for the FkhP motif, although their study included non-physiological pH values as low as pH 5 and as high as pH 9. Using FOXP1, Medina and colleagues ^89^ show that the protonation state of a less conserved histidine exclusive to the FOXM/O/P subfamilies modulates domain swapping stability to regulate DNA-binding affinity. Collectively, these findings and our current study strongly support a pH-regulated binding of FOX family transcription factors to an FkhP motif with higher affinity binding at lower pH.

Our corollary prediction, that a deprotonated histidine at higher pH favors binding to adenine by acting as a hydrogen bond acceptor, is suggested by our SELEX-seq and RNA-seq data. Both SELEX and RNA-seq as unbiased approaches reveal pH-dependent binding preferences with lower pH favoring histidine-thymine hydrogen bonds and higher pH favoring histidine binding AGC rich sequences. However, we could not directly confirm *in vitro* higher affinity binding at higher pH to GACGC identified by SELEX-seq that resembles a reported FOX FHL-N consensus motif ^59^. However, SELEX-seq with FOXC2 identified distinct DNA sequences that bind at pH 7.0 compared with pH 7.8. Two possibilities could account for our inability to confirm *in vitro* binding to the motifs identified at higher pH. First, PCR amplification steps in SELEX-seq may overrepresent overall binding affinity of FOXC2. Second, the short 5 base pair sequences or 2x repeats might not been sufficient for FOXC2 binding without flanking nucleotides present in the SELEX library.

Additionally, our findings from RNA-seq confirm that FOXC2 regulates the expression of different genes at higher compared with lower pHi. Several of the top pHi- and FOXC2-His122- dependent genes at higher pH increase expression of tumor promoting genes while simultaneously attenuating expression of tumor suppressive genes that are enriched at low pHi. These results are consistent with previous findings that increased pHi promotes cancer cell behaviours including cell cycle progression, epithelial to mesenchymal transition, migration, and invasion ^3,13–15,52,58,90,91^. These results also suggest a possible mechanism for how pHi directly regulates gene expression through pH-sensing of transcription factors and further highlights pHi as a signaling mechanism regulating protein electrostatics for changes in cell behaviours. Further, our RNA-seq results also underscore how relatively small changes in pHi can lead to large scale changes in a gene expression as well as predicted DNA motif-binding preferences. These findings are consistent with our previous reports on pH sensing by endogenous proteins with small changes in the cellular pH range conferring cell behaviours, including stability of β-catenin by pH-regulated binding to the E3 ligase β-TrCP regulating epithelial cell integrity ^30^, activity of the focal adhesion kinase FAK through in-cis conformational changes conferring cell substrate adhesion dynamics ^7^, and the affinity of talin for binding F-actin in controlling cell migration ^6^.

The concept of histidine titration conferring nucleotide-binding selectivity can be applied to a broader scope. First, although our study focused exclusively on FOX family members, pH-dependent binding of transcription factors to DNA may be a relevant to transcription factors in other families such as KLF, SOX, and MITF/MYC/MAX that contain a conserved histidine in the DNA binding domain, which in available structures forms a hydrogen bond with nucleotides. For the SOX family, Sry and SOX30 are the only members that lack a nucleotide-binding histidine, and instead contain a cognate arginine and asparagine, respectively. Future studies could use DBD swapping with Sry and SOX30, which cannot be applied to FOX proteins that contain an invariant histidine. Second, the concept can be applied to cell behaviours regulated by pHi dynamics that include changes in gene expression, most notably stem cell differentiation and lineage specification ^10–12^. Third, we propose that histidine titration conferring nucleotide-binding selectivity could in part determine how different co-expressed members of transcription factor families with highly conserved DNA-binding domains recognize distinct DNA sequences and are used reiteratively to regulate diverse target genes and disparate cell behaviours, which are unresolved questions for understanding developmental processes. Finally, beyond transcription factors, some RNA binding proteins contain a critical nucleotide-binding histidine ^92^.

Based on our current findings and applications for a broader scope, we propose that pH dynamics is an understudied mechanism regulating nucleotide binding by proteins with functionally critical histidine residues. Moreover, we highlight how pHi regulated electrostatics can affect protein functions for diverse cell behaviours for a more complete understanding of both development and disease.

## Acknowledgements

This work was supported by the National Institutes of Health grants R01 CA197855 to D.L.B. and F31 GM142284 to K.P.K., as well as a National Science Foundation grant NSF 2203629 and a UCSF Sandler Program for Breakthrough Biomedical Research award to D.L.B. and M.P.J. We also thank Connie Phuong from the Barber laboratory and the members of the Mark Anderson laboratory at UCSF for technical assistance with generating recombinant protein described in this work.

---

**Supplementary Figure S1.** Histidine in the DBD of transcription factors in different families. A-D. Representative transcription factors from families with conserved histidine (magenta) forming a hydrogen bond with DNA nucleotides, including FOXC2-His122 binding to thymine (PDB:6AKO) (A), KLF4-His474 binding to adenine (PDB: 2WBU) (B), SOX4-His29 binding to guanine (PDB: 3U2B) (C), and MITF-His209 binding to guanine (PDB: 4ATK) (D). E-G. Select transcription factors in other families shown to have a hydrogen bond between histidine (magenta) and a DNA nucleotide, including ETV6-His396 binding to thymine (PDB: 4MHG) (E), STAT6-H415 binding to guanine (PDB:4Y5W) (F), and ARNT-His94 binding to guanine (PDB: 4ZPK) (G).

**Supplementary Figure S2.** Purified GST fusion proteins of FOXC2, FOXM1, and FOXN1 DBD. A-C. Coomasie stained gels of purified GST fusions of the DBDs of FOXC2 (A), FOXM1 (B), and FOXN1 (C).

**Supplementary Figure S3.** Measured and predicted binding of FOXM1-H287K to an FkhP sequence. A, B. Binding of recombinant DBD of FOXM1-H287K, including binding curves at the indicated pH values, determined by fluorescence anisotropy (A) and association constants calculated from binding curves (B). Data are means ± s.e.m. of three separate measurements with two independent protein preparations. C. CpHMD portion of simulation in which FOXM1 DBD HIS residues are double protonated (HIP), protonated at delta nitrogen (HID), protonated at epsilon nitrogen (HIE) are predicted to be unbound (circles) or bound FkhP (slashes). All DBD HIS residues are labeled His269 (blue), His275 (cyan), His287 (orange), His292 (yellow), His311 (green).

**Supplementary Figure S4.** Loss of NHE1 activity and expression of recombinant FOXC2 in NHE1-KO MDA-MB-436 cells. A. Confocal images of control and EIPA treated MDA-MB-436 cells loaded with SNARF. Nuclei are indicated by dashed circles. B. The pHi recovery in a nominally-HCO_3_-free Hepes buffer from an NH_4_Cl-induced acid load, an index of NHE1-dependent H+ extrusion, in the indicated MDA-MB-436 cells in the absence and presence of EIPA and with NHE1-KO loaded with the pH-sensitive dye BCECF. B. FOXC2 expression from luciferase assay lysates in controls and NHE1-KO MDA-MB-436 cells.

