## Supplementary Figures for "A role for pH dynamics regulating transcription factor DNA binding selectivity"

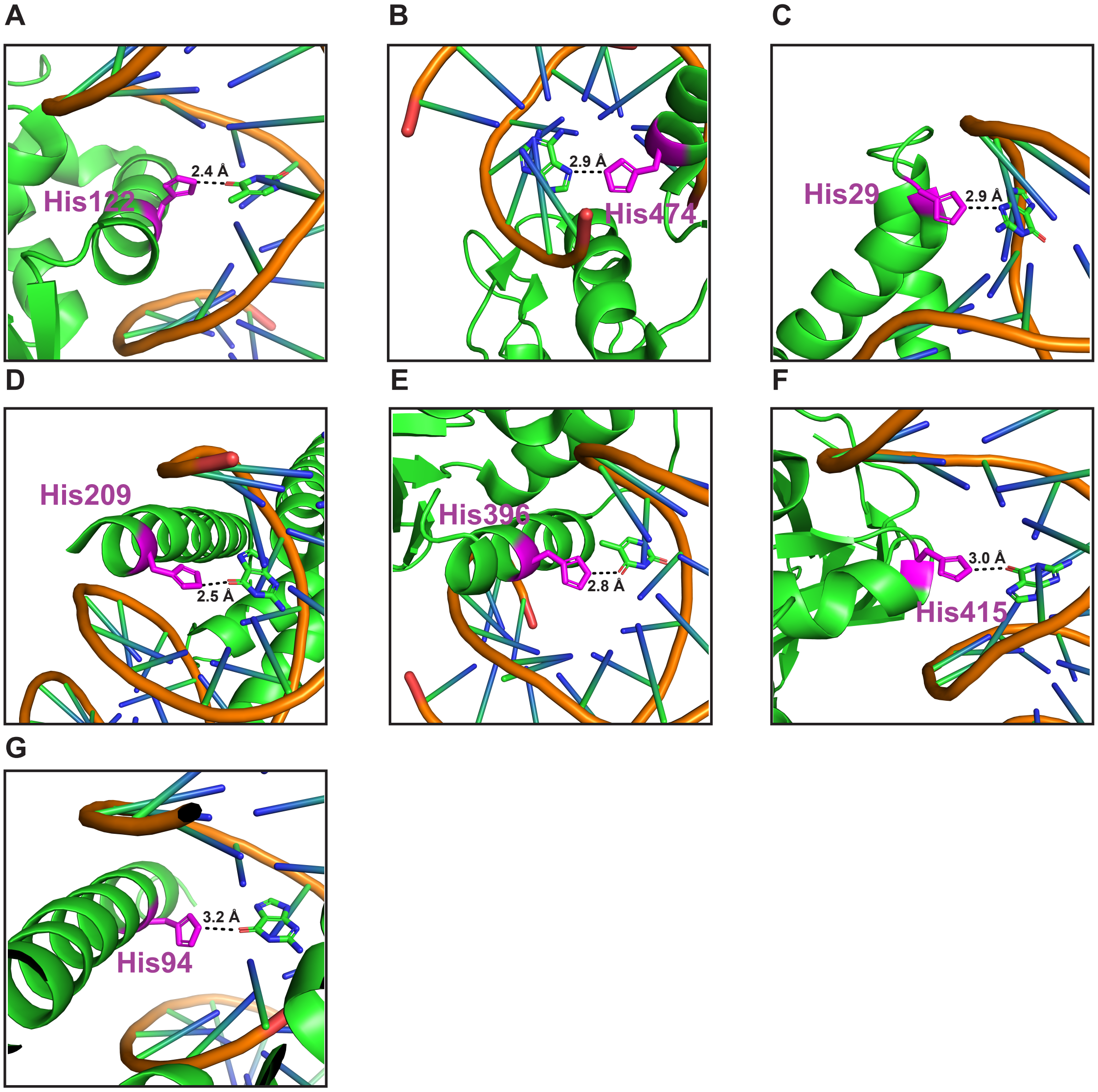


**Supplementary Figure 1. Kisor et al.**

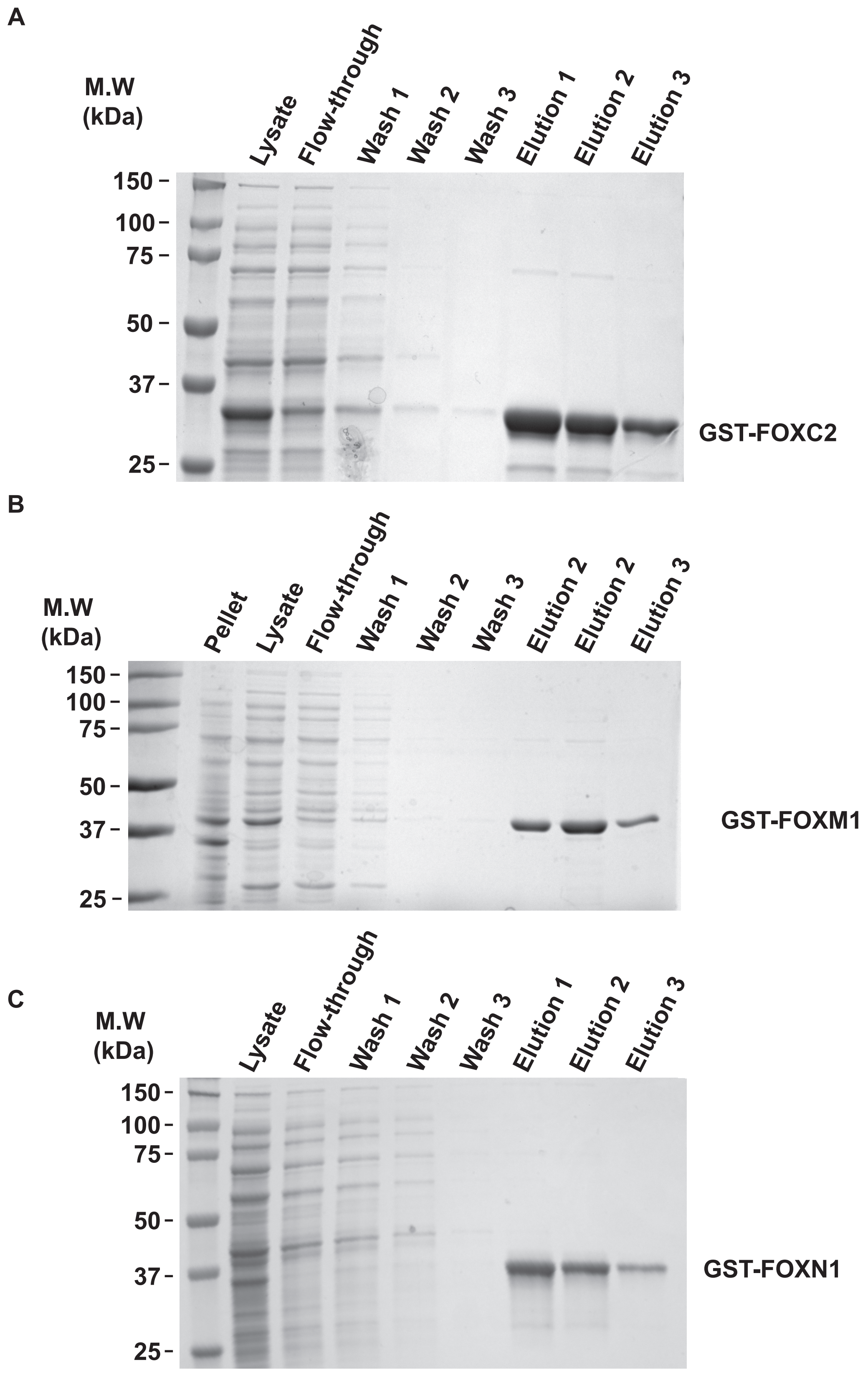


**Supplementary Figure 2. Kisor et al.**

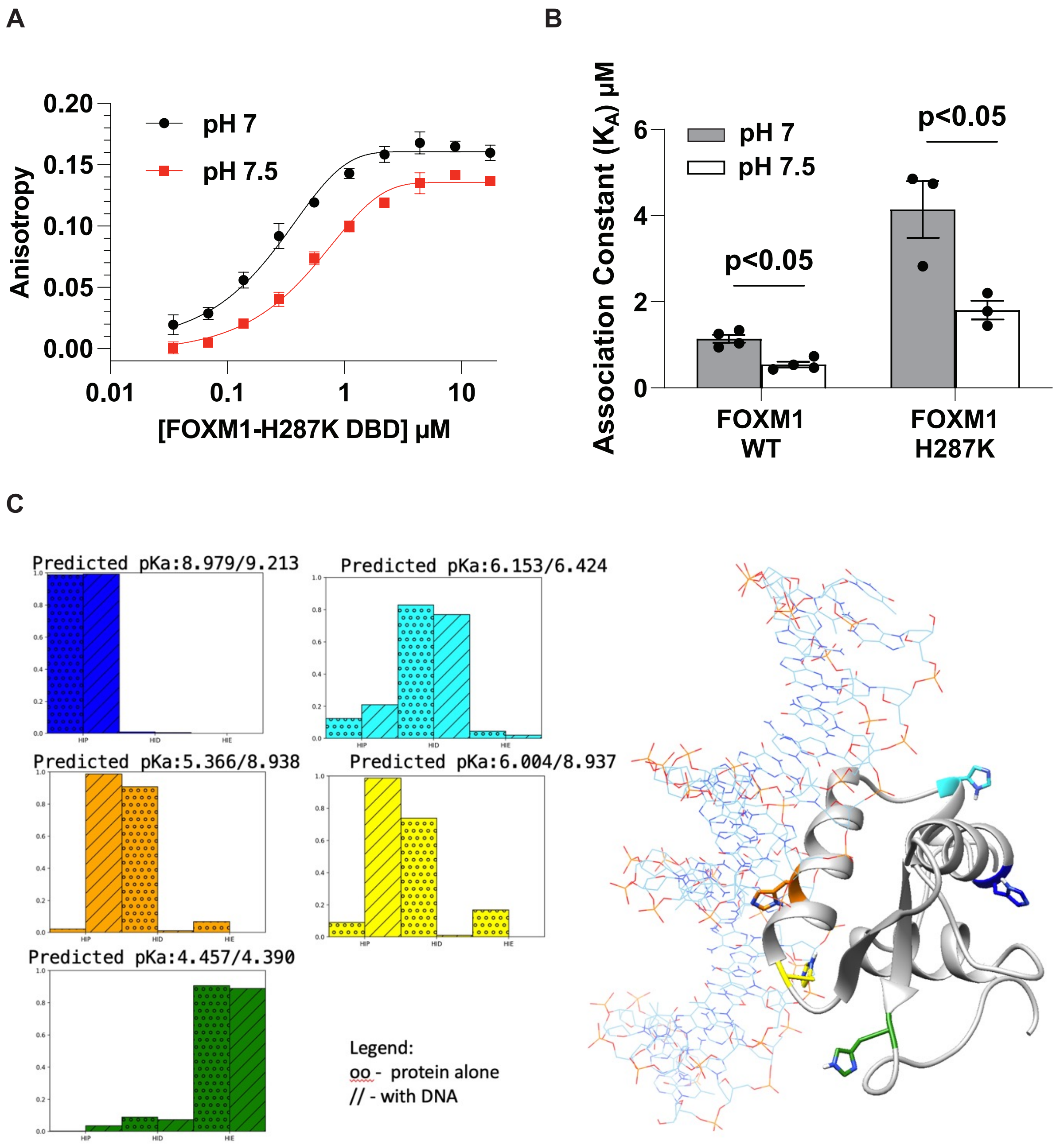


**Supplementary Figure 3. Kisor et al.**

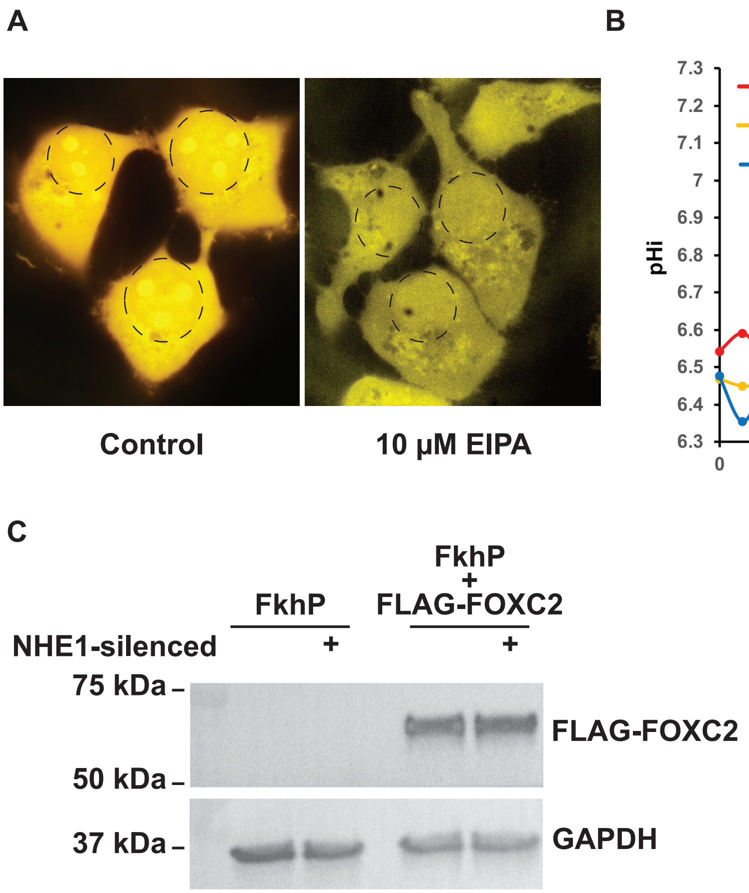


**Supplementary Figure 4. Kisor et al.**
